# The Precise Basecalling of Short-Read Nanopore Sequencing

**DOI:** 10.1101/2024.09.12.612746

**Authors:** Ziyuan Wang, Mei-Juan Tu, Chengcheng Song, Ziyang Liu, Katherine K. Wang, Shuibing Chen, Ai-Ming Yu, Hongxu Ding

## Abstract

The nanopore sequencing of short sequences, whose lengths are typically less than 0.3kb therefore comparable with Illumina sequencing techniques, has recently gained wide attention. Here, we design a scheme for training nanopore basecallers that are specialized for short biomolecules. With bioengineered RNA (BioRNA) molecules as examples, we demonstrate the superior accuracy of basecallers trained by our scheme.

## INTRODUCTION

Nanopore sequencing is known for its capability of long-read sequencing: with the latest Ultra-Long DNA Sequencing Kit V14 provided by Oxford Nanopore Technologies (ONT), a >50kb N50 could be achieved. This capability greatly facilitated biological applications such as *de novo* genome assembly and RNA splicing isoforms determination, which are challenging for traditional Next Generation Sequencing-based approaches^1^. Conversely, the nanopore sequencing of short molecules, which are typically shorter than 0.3kb, has recently gained wide attention. For instance, previous studies have proved the feasibility of sequencing tRNAs, whose lengths are 76-90 nucleotides^2-5^. The possibility of profiling ever shorter biomolecules, such as microRNAs (21-25 nucleotides) were also explored^6^. Furthermore, a general method for sequencing short non-coding RNAs, including tRNAs and microRNAs, were developed^7^. While sequencing biomolecules promises substantial scientific insights, sequencing artificially synthesized oligos could facilitate the training of bioinformatic models. For instance, a group of chemically-modified, 46 nucleotide-length short-oligos were used as training data for detecting 8-hydroxyguanine^8^. Noticeably, the 0.3kb read length is comparable with Illumina techniques, which typically produce reads ranging from 0.05 to 0.3kb. Albeit the growing interest, bioinformatic methods tailored to short-read nanopore sequencing remain lacking. To close such a knowledge gap, in this study, we develop a scheme for training basecallers that could accurately interpret short nanopore sequencing readouts.

## RESULTS

### The nanopore sequencing of BioRNAs

We adopted BioRNAs as models to study the short-read basecalling. BioRNAs are RNA interference (RNAi) agents made *in vivo*, and some have demonstrated therapeutic effects across a range of diseases^9^. The BioRNA complex is engineered from specific human tRNA, with the RNAi precursor (pre-miRNA) replacing the anticodon sequence as shown in Figure 1A. Upon the intracellular uptake, miRNA processing proteins cleave the pre-miRNA, further releasing the functional RNAi molecule to exert pharmacological actions. We prepared BioRNA nanopore sequencing libraries following the Nano-tRNAseq protocol^4^ (Figure 1B, see METHODS). Specifically, we sequenced four BioRNAs including two molecules whose anticodon was replaced by a Sephadex aptamer (BioRNA^Ser^ and BioRNA^Leu^), as well as two therapeutic candidates whose anticodon was replaced by a pre-miRNA-34a sequence (BioRNA^Ser^/miR-34a and BioRNA^Leu^/miR-34a, see METHODS). The sequences, as well as the multiple sequence alignment (MSA) result, of the four BioRNA molecules were presented in Figure 1C.

**Figure 1.**
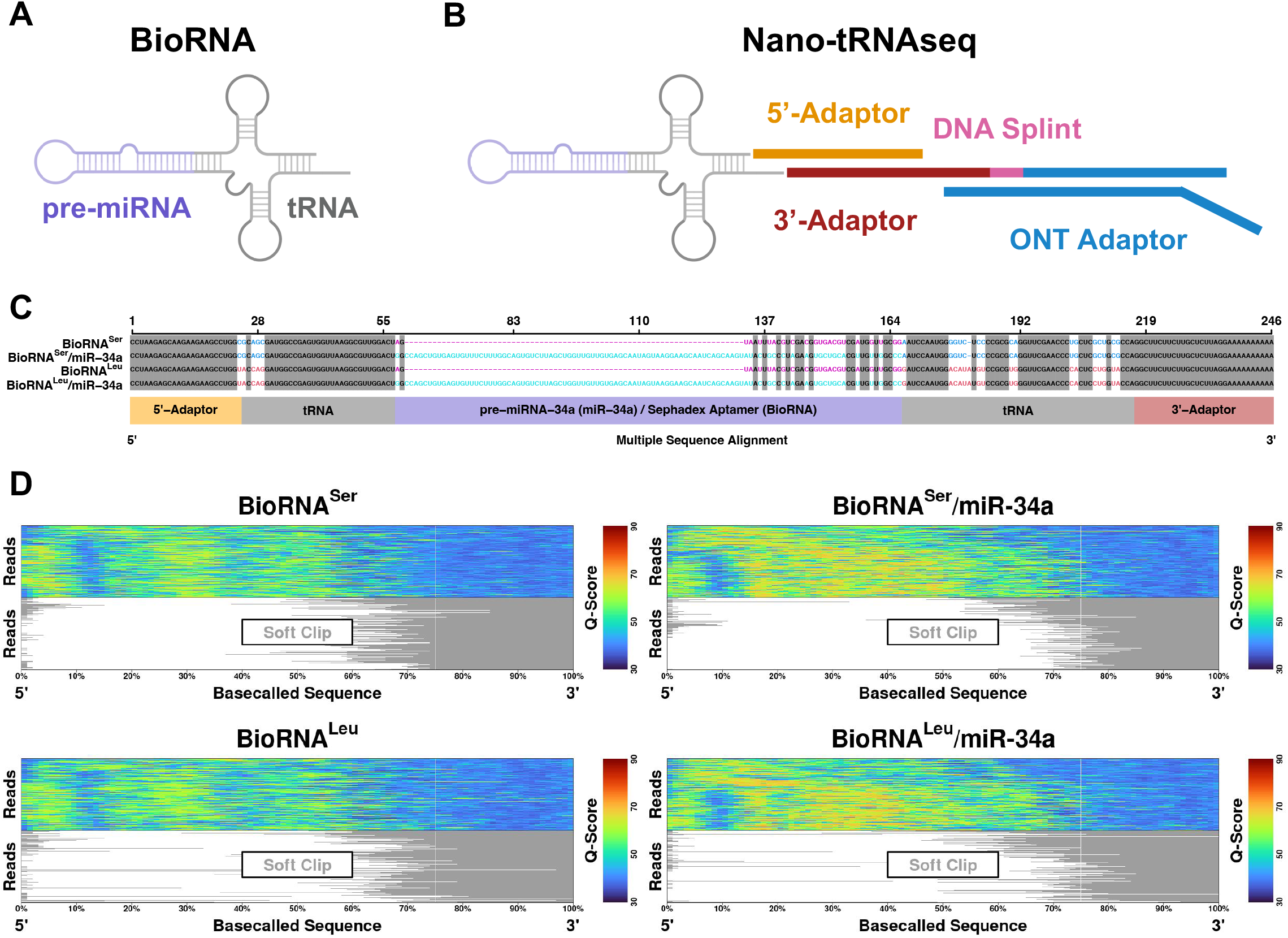
Compromised basecalling accuracy on 3’ and 5’-ends of short nanopore sequencing readouts. (A) The BioRNA therapeutic RNA interference (RNAi) molecule. Preparing BioRNA nanopore sequencing libraries using the Nano-tRNAseq strategy. The multiple sequence alignment of BioRNA^Ser^, BioRNA^Ser^/miR-34a, BioRNA^Leu^, and BioRNA^Leu^/miR-34a Nano-tRNAseq nanopore sequencing libraries. (D) The visualization of Q-scores and soft-clipped regions along individual basecalled sequences. Sequences were normalized by lengths and visualized by length-percentages.

### Compromised basecalling accuracy on 3’ and 5’-ends

We first quantified the Bonito (the state-of-the-art basecaller released by ONT) basecalling accuracy on BioRNAs. We noticed decreased Q-scores, which indicate a less confident basecalling, on both 3’ and 5’-ends. In particular, average Q-scores within the last ∼50 nucleotides in the 3’-end and the first ∼4 nucleotides in the 5’-end decreased to ∼40. In contrast, the average Q-score of the BioRNA gene-body remained ∼60. We further found that the less confident 3’ and 5’-regions coincided with unalignable 3’ and 5’-end soft-clips. Considering the lengths of BioRNAs are relatively short (∼200 nucleotides), >40% of basecalled nucleotides cannot be properly aligned (Figure 1D, see METHODS). We therefore concluded widespread 3’ and 5’-artifacts as the major bioinformatic challenge in short-read nanopore sequencing.

### A basecaller training scheme to polish error-prone 3’ and 5’-ends

Our recent study demonstrated that compromised basecalling can be polished via an iterative workflow^10^. Our workflow builds upon the community consensus that Bonito can achieve acceptable accuracy for generic basecalling tasks. We then polish the yielded sketch sequences by aligning them to the ground-truth reference. We next take polished sequences, and their nanopore sequencing readouts, as training data to update Bonito. To make our workflow more effective in polishing 3’ and 5’-ends, we further develop a 3-step sampling strategy for preparing training data. Specifically, we sample reads that are mapped to the 5’-end, the entire molecule and the 3’-end subsequently. Our strategy guarantees the candidate sequence to be evenly covered by training reads, and therefore promises to improve the basecalling of 3’ and 5’-ends (Figure 2A and B, see METHODS).

**Figure 2.**
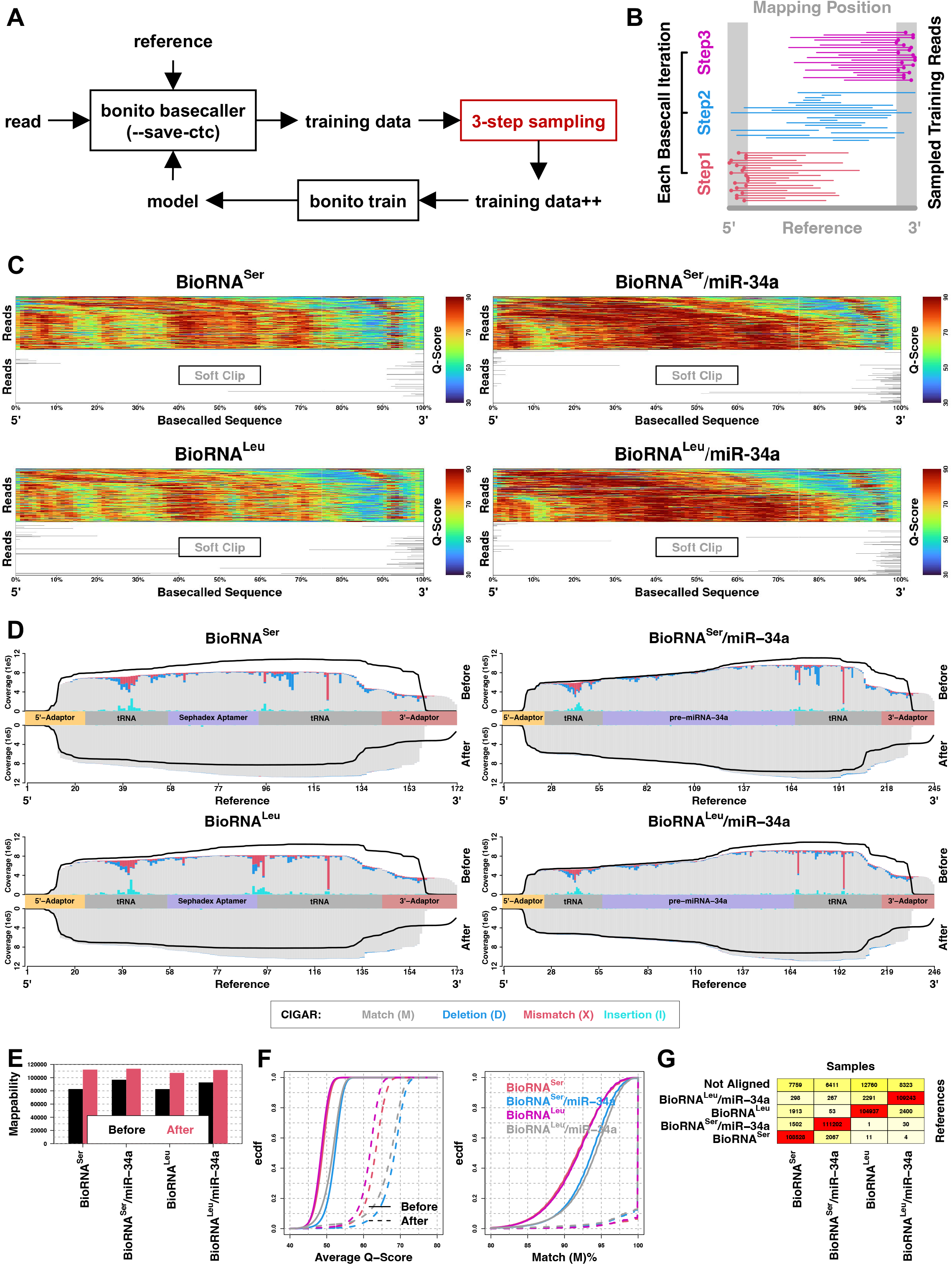
Polishing short RNA sequences via iterative basecalling. (A) The iterative basecalling workflow. (B) The 3-step sampling strategy for preparing training data during iterative basecalling. Specifically, first and third steps prioritize reads that are mapped to 5’ and 3’-ends respectively, the second step has no selection on read mapping position. This strategy guarantees the entire candidate molecule could be covered by high-quality training data, therefore improving the basecalling of 3’ and 5’-ends. (C) The visualization of Q-scores and soft-clipped regions along individual basecalled sequences. Sequences were normalized by lengths and visualized by length-percentages. (D) Alignment results of BioRNAs. The per-site coverage and CIGAR status were visualized. Before and After denote the vanilla Bonito and iterative basecalling, respectively. Inside each subplot, the black curve mirrors the coverage of its counterpart. For each BioRNA species, a total of 120,000 reads were analyzed. (E) Mappabilities, which quantify total numbers of aligned reads, of BioRNAs. (F) Distributions of the per-read average Q-score and CIGAR Match percentage. Ecdf, the empirical cumulative distribution function. (G) Alignment status of BioRNAs after the iterative basecalling.

We polished BioRNA 3’ and 5’-ends with the iterative training scheme. We found greatly improved 3’ and 5’-basecalling confidence with average Q-scores increased to ∼60 from ∼40. Besides 3’ and 5’-ends, basecalling confidence of the BioRNA gene-body was also improved, with average Q-scores increased to ∼85 from ∼65 (Figure 2C). Consequently, basecalling artifacts quantified by alignment anomalies, including 3’ and 5’-end soft-clips (Figure 2C) and mismatches, deletions and insertions inside the gene-body (Figure 2D), were greatly resolved (see METHODS). Such polished basecalling further promoted the mappability among all the BioRNA groups: we noticed an average ∼20% increase in the number of aligned reads. We also concluded improvements on the per-read basecalling confidence and accuracy in Figure 2F. We finally demonstrated that the 3-step sampling strategy outperformed the default iterative basecaller training setup, by recovering more mis-basecalled nucleotides on, in particular, the 5’-end (Figure S1).

### Distinguishing homologous molecules with polished basecalling

We further tested if iterative basecalling results are accurate enough to differentiate highly-similar BioRNA sequences. We found that 87% BioRNA^Ser^ and BioRNA^Leu^, and 91% BioRNA^Ser^/miR-34a and BioRNA^Leu^/miR-34a sequences are matched. Additionally, BioRNA^Ser^ and BioRNA^Leu^ share the same carrier tRNA molecule with BioRNA^Ser^/miR-34a and BioRNA^Leu^/miR-34a, respectively (Figure 1C). We challenged our iterative basecalling with such homologous molecules, and observed superior basecalling accuracy. Specifically, an average of 90% reads could be correctly aligned to corresponding reference, e.g. BioRNA^Ser^ reads to the BioRNA^Ser^ sequence. However, there were still an average of 7.3% reads that cannot be assigned to any references, as well as an average of 2.7% misaligned reads (Figure 2F, see METHODS). We noticed unalignments to be sequencing readouts with longer dwell time but shorter sequence lengths (Figure S2), which were likely produced from blocked nanopores. We noticed misalignments between BioRNA^Ser^ and BioRNA^Ser^/miR-34a, and BioRNA^Leu^ and BioRNA^Leu^/miR-34a are in general truncations that mapped to the 3’-end, where the universal Nano-tRNAseq 3’-adaptor locates. Therefore, it is extremely difficult to align these truncated readouts. We further noticed misalignments between BioRNA^Ser^ and BioRNA^Leu^, and BioRNA^Ser^/miR-34a and BioRNA^Leu^/miR-34a to be readouts with low MAPQ scores, which are most likely to be alignment artifacts (Figure S3).

## DISCUSSION

Short-read sequencing emerges as the new research focus in the nanopore sequencing community. Here, we report a scheme for training basecallers to precisely interpret short nanopore readouts. Our scheme resolves the widespread 3’ and 5’-basecalling artifacts, which can affect >50 RNA nucleotides (>15% length of a 0.3kb molecule) therefore may significantly compromise downstream bioinformatic analyses, through balancing training reads to cover both 3’ and 5’-ends. We demonstrated the efficacy of this scheme, by the accurate basecalling of short, homologous BioRNA therapeutic RNAi species. To further extend our success, we formulated the following two future directions.

### Extending the 3-step sampling strategy for training long-read basecallers

We also found widespread 3’ and 5’-basecalling artifacts among long-read nanopore sequencing readouts. As an example, we sequenced ∼2kb RNA control oligos (see METHODS) and noticed decreased Q-scores together with excessive soft-clips on 3’ and 5’-ends (Figure S4A). Such commonly-existing 3’ and 5’-basecalling artifacts might substantially impede the applicability of nanopore sequencing in fundamental and translational studies. It has been demonstrated that mRNA 3’ and 5’-Untranslated Regions (UTRs) play critical roles in eukaryotic translation initiation and regulation^11^. Optimized UTRs have therefore been adopted to augment the *in vivo* translation of therapeutic mRNAs^12^. The 3-step sampling strategy, on the other hand, can largely reduce such artifacts further producing accurate basecalling results (Figure S4B). Taken together, we proposed the 3-step sampling as a generic strategy for training precise nanopore basecallers.

### Introducing 3’ and 5’-adaptors to protect the basecalling of target sequences

Our 3-step sampling strategy failed to properly basecall the first ∼10 5’-positions and the last ∼15 3’-positions among all the four BioRNA types (Figure 2D). This is because for these end-regions, the initial basecalling by the vanilla Bonito model generally cannot produce enough high-quality training reads. Thus, even with the 3-step sampling strategy, the far ends of BioRNAs still cannot be precisely represented. Fortunately, compromised 3’ and 5’-basecalling only influenced Nano-tRNAseq adaptor regions. BioRNA gene bodies, on the other hand, were properly characterized with negligible artifacts. We thus concluded the indispensability of 3’ and 5’-protective adaptors in basecalling target molecules.

## METHODS

### The Production and Purification of BioRNAs

The design and production of bioengineered human seryl-tRNA (TGA) and leucyl-tRNA (TAA) (BioRNA^Ser^ and BioRNA^Leu^), and tRNA-fused human pre-miR-34a molecules (BioRNA^Ser^/miR-34a and BioRNA^Leu^/miR-34a) was described in our previous studies^13-15^. In brief, sequences of human tRNAs and pre-miR-34a were obtained from GtRNAdb (https://gtrnadb.ucsc.edu/) and miRbase (https://www.mirbase.org/), respectively. For BioRNA^Ser^ and BioRNA^Leu^, a Sephadex aptamer was inserted to substitute the anticodon sequence of the corresponding tRNA. For BioRNA^Ser^/miR-34a and BioRNA^Leu^/miR-34a, the optimized pre-miR-34a sequence^16^ was inserted into the anticodon region of the corresponding tRNA. Coding sequences of BioRNAs were summarized in Table S1.

These sequences were amplified using polymerase chain reaction (PCR), and the yielded amplicons were infused in the pBSTNAV vector with the Takara In-Fusion® HD Cloning kit to construct RNA expression plasmids. After being verified with the DNA sequencing, plasmids were transformed into *E. coli* HST08 competent cells to express and accumulate BioRNAs by overnight fermentation on a large scale (∼500 mL). The total RNA was isolated from the overnight culture by the phenol extraction, and subsequently loaded and separated with the Bio-Rad ENrich-Q 10×100 anion exchange column integrated into the Bio-Rad NGC Quest 10 Plus chromatography system for purifying target BioRNAs. The total RNA was eluted with the gradient method, which consist of buffer A (10 mM monosodium phosphate, pH 7.0) and buffer B (buffer A + 1 M sodium chloride, pH 7.0) at a constant flow rate of 2 mL/min. To purify BioRNAs, the column was initially equilibrated by buffer A for 6.7 min, followed by elution with 55% buffer B for 4.8 min, then 55-70% buffer B for 40 min (BioRNA^Ser^), or 55-75% buffer B for 30 min (BioRNA^Leu^ and miR-34a BioRNAs). The column was subsequently washed with 100% buffer B for 10 min, with an additional re-equilibration using buffer A for 10 min. Fractions of target RNA peaks were collected during the total RNA elution, and loaded onto urea PAGE gels to verify sizes and homogeneity. Pure fractions were combined, desalted and concentrated with Amicon Ultra-2mL centrifugal filters (30 k) to obtain ready-to-use pure BioRNAs. The BioRNA purity was determined by urea PAGE gel analyses, and quantified by high-performance liquid chromatography (HPLC) as previously reported^14,17^. BioRNA products with high homogeneity (> 98%) were used for nanopore sequencing.

### The Synthesis of Nano-tRNAseq Adaptors

The chemically synthesized Nano-tRNAseq^4^ adapters were ordered from Integrated DNA Technologies. Specifically, for nano-tRNAseq 5’ RNA splint adapters, the first four nucleotides at the 3’ terminal were designed as rUrGrGrC and rUrGrGrU, which are complementary to seryl-tRNA and leucyl-tRNA, respectively. Other Nano-tRNAseq adapters, including 3’ RNA:DNA splint adapters and ONT reverse transcription adapters (RTAs, oligo A and oligo B) were synthesized as reported in the original study. Sequences of Nano-tRNAseq adapters were listed in Table S1.

### The Nanopore Sequencing of BioRNAs

BioRNA nanopore sequencing libraries were made first following the nano-tRNAseq protocol^9^, then using the Oxford Nanopore Technologies Direct RNA Sequencing Kit (SQK-RNA004) following manufacturer’s instructions. Specifically:

#### Adapter Annealing

5’ RNA splint adapters and equal moles (∼108 pmol) of 3’ RNA:DNA splint adapters were mixed with a dilution buffer consisting of Tris-HCl (10 mM, pH 7.5), NaCl (500 mM), and 1 μL of RNasin Ribonuclease Inhibitor to a final concentration of ∼50 ng/μL for each adapter in a total volume of 20 μL. The mixture was incubated at 75 °C for 15 seconds, further cooled to 25 °C at a rate of 0.1 °C/s (a total of 500 seconds) to anneal splint adapters. RTA oligo As and oligo Bs (∼82 pmol for each, ∼200 ng in total) were annealed under the same condition as splint adapters.

#### Ligation 1

1 μg of BioRNA^Ser^, BioRNA^Leu^, BioRNA^Ser^/miR-34a or BioRNA^Leu^/miR-34a was ligated to pre-annealed splint adapters (RNA:adapters = 1.2:1, molar ratio) in a 20 μL reaction system containing T4 RNA ligase 2, 1 × NEB Reaction Buffer, 10% PEG 8000, 400 μM ATP, 1 μL RNasin Ribonuclease Inhibitor. The reaction was conducted at room temperature for 1 hour. Ligation products were purified with 1.8x volume of well-mixed, room temperature AMPure RNAClean XP beads following manufacturer’s instructions. Concentrations of purified samples were determined using the Tecan plate reader, and the quality of ligated RNAs was evaluated by the urea PAGE gel.

#### Ligation 2 and reverse transcription (RT)

Purified “Ligation 1” products (400 ng, for two sequencing replicates) were then ligated to pre-annealed RTA adapters (RNA:adapters = 1:2, molar ratio) with the NEB T4 DNA Ligase in a 30 μL reaction buffer constituted by 6 μL of NEBNext Quick Ligation Reaction Buffer, 1 μL of RNasin Ribonuclease Inhibitor and RNase-free water by incubating at room temperature for 30 min. Subsequently, the ligation product was mixed with 4 μL of dNTPs (10 mM) and 26 μL of RNase-free water to pre-denature the RNA at 65 °C for 5 min and then cooled down on ice for 2 min. A mixture of 4 μL Maxima H Minus Reverse Transcriptase, 16 μL of Maxima H Minus Reverse Transcriptase Buffer, and 2μL of RNasin Ribonuclease Inhibitor (80U) were added to the pre-treated RNA sample to perform RT by incubation at 60 °C for 1 hour, 85 °C for 5 min, then cooling to 4 °C. AMPure RNAClean XP beads were then added to purify the RT products according to the protocol. The quantity and quality of the purified samples were determined by the Tecan plate reader and urea PAGE gels, respectively.

#### Ligation 3 and final library

Following purification, 40 μL (two replicates) of RT products were ligated to 12 μL of RNA Ligation Adapter (RLA, provided by the SQK-RNA004 kit) in an 80 μL reaction system, supplemented with 16 μL of NEBNext Quick Ligation Reaction Buffer and 6 μL of T4 DNA Ligase for a 30-min incubation at room temperature. The ligation products were purified using AMPure RNAClean XP beads following the protocol provided by the SQK-RNA004 kit. Afterwards, MinION flow cells (RNA chemistry) were primed following manufacturer’s instructions. Meanwhile, Elution Buffer (provided by the SQK-RNA004 kit) re-suspended libraries were gently mixed with 37.5 μL of Sequencing Buffer (provided by the SQK-RNA004 kit) and 25.5 μL of Library Solution (provided by the SQK-RNA004 kit) to obtain final libraries. A total volume of 75 μL final library was loaded to each flow cell for sequencing.

### The Design, Production and Nanopore Sequencing of Curlcake RNA Oligos

We designed oligo sequences following procedures reported in ^18^ using the CURLCAKE software (http://cb.csail.mit.edu/cb/curlcake/). Specifically, we covered all possible RNA 5mers (1,024 in total, median occurrence as 10), and avoided secondary structures. We splitted the CURLCAKE sequence (12.5 kb) as four “curlcakes” of ∼2.5 kb for synthesis purposes. For each RNA curlcake, we first added the strong T7 promoter to the 5’ end. We then added EcoRV sites to both 3’ and 5’ ends, and removed all the internal EcoRV and BamHI sites. RNA sequences were summarized in Table S2. We synthesized and cloned all the RNA curlcakes into the pUC57 vector using blunt EcoRV and HindIII, with the service of GenScript Biotech Corporation.

Curlcake RNA oligos were produced by *in vitro* transcription (IVT). Engineered pUC57 plasmids were digested by EcoRV and BamHI restriction enzymes for at least two hours at 37 ^°^C, and analyzed via agarose gel electrophoresis. The digested DNA was purified by a PCR purification kit. Nanodrop was used to measure the concentration of extracted DNA prior to IVT. The Ampliscribe™ T7-Flash™ Transcription Kit was used to make IVT RNAs following manufacturer’s instructions. DNAse I was added to the IVT reaction system after incubation for 4 hours at 42 ^°^C to eliminate the residual template DNA. IVT RNAs were purified using the RNeasy Mini Kit following manufacturer’s instructions. NEB vaccinia capping enzyme was used for the 5’ capping of purified IVT RNAs, with an incubation for 30 min at 37 ^°^C. Following purification with RNAClean XP Beads, the capped IVT RNAs were subjected to polyadenylation tailing. Concentration of capped and polyA-tailed IVT RNAs was determined by Qubit Fluorometric Quantitation.

RNA nanopore sequencing libraries were built using the ONT Direct RNA Sequencing Kit (SQK-RNA004) following protocol version DRS_9195_v4_revC_20Sep2023-minion as per manufacturer’s instructions. Briefly, 2 μg of capped and polyA-tailed IVT RNA was subjected to adapter ligation using the NEB T4 DNA Ligase, following reverse transcription using the SuperScript III Reverse Transcriptase. After purification using RNAClean XP Beads, yielded RNA:DNA hybrids were ligated to RNA adapters using the NEB T4 DNA Ligase. The concentration of the yielded RNA library was determined by the Qubit fluorometer. The RNA library was mixed with RNA Running Buffer prior to sequencing on a primed MinION flow cell. The flow cell chemistry is RNA, and the sequencer used is MinION.

## Data Availability

The BioRNA nanopore sequencing data was deposited at NCBI under the BioProject PRJNA1155679 (reviewer link: https://dataview.ncbi.nlm.nih.gov/object/PRJNA1155679?reviewer=tmp2haiuk3d28bd7q25p15ula0). The curlcake nanopore sequencing data was deposited at NCBI under the BioProject PRJNA1154950 (https://dataview.ncbi.nlm.nih.gov/object/PRJNA1154950?reviewer=koot1fburqsrjg1d0t426sk92s).

## Code Availability

The Bonito iterative basecalling framework can be found at: https://github.com/wangziyuan66/iterative-labeling-toolkit-bonito.

## ACKNOWLEDGEMENTS

We thank the Cornell University Center for Advanced Computing team, University of Arizona High Performance Computing team and the College of Pharmacy Information Technology Group for their support. H.D. is supported by the University of Arizona Health Sciences Career Development Award. A.-M.Y. is supported by the National Institute of General Medical Sciences [R35GM140835] and National Cancer Institute [R01CA225958 and R01CA253230], National Institutes of Health (NIH).

## AUTHOR CONTRIBUTIONS

Z.W. and H.D. conceived the idea. Z.W. and Z.L. performed the analysis. M.-J.T., C.S. and K.K.W. performed the experiment. S.C., A.-M.Y. and H.D. supervised the project. All authors wrote the manuscript.

