## Supplementary for "The Precise Basecalling of Short-Read Nanopore Sequencing"

### Supplementary Information

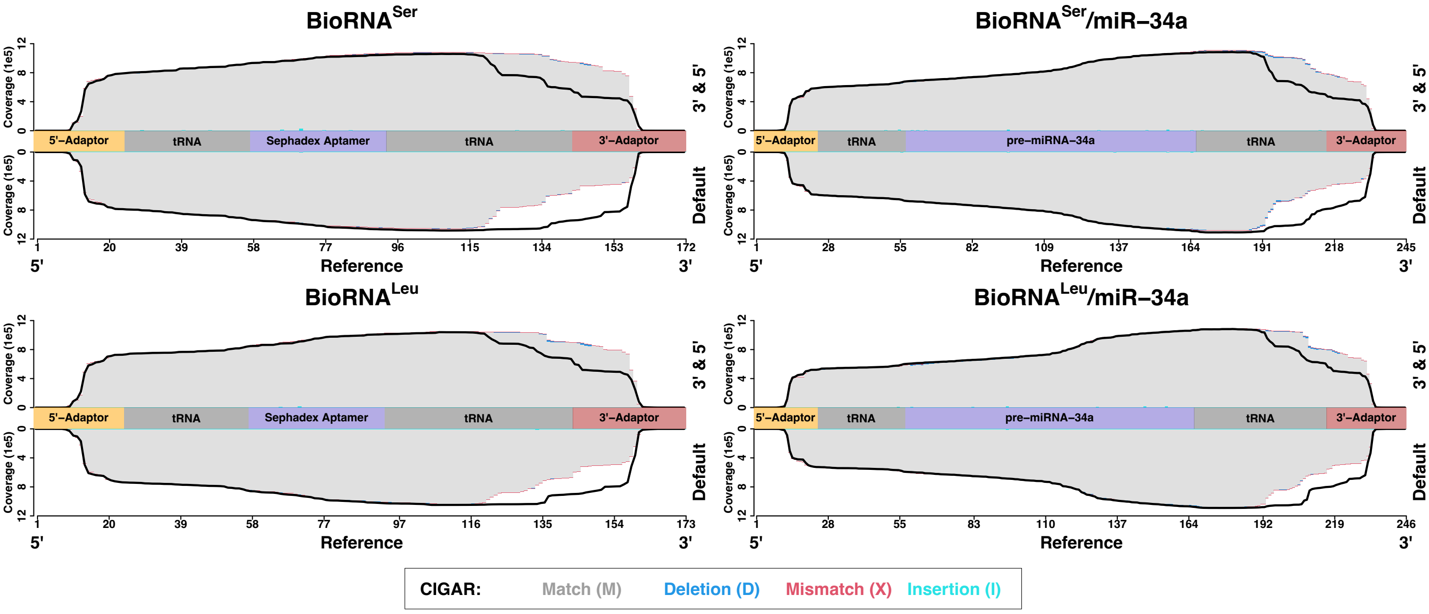

BioRNA^Ser
BioRNA^Ser/miR–34a
BioRNA^Leu
BioRNA^Leu/miR–34a
Coverage (1e5)
3' & 5'
Default
5'–Adaptor
tRNA
Sephadex Aptamer
pre–miRNA–34a
3'–Adaptor
Reference
5'
3'
1 20 39 58 77 96 115 134 153 172
1 28 55 82 109 137 164 191 218 245
1 20 39 58 77 97 116 135 154 173
1 28 55 83 110 137 164 192 219 246
CIGAR:
Match (M)
Deletion (D)
Mismatch (X)
Insertion (I)

**Figure S1. Basecalling comparison between the 3-step sampling strategy and the default iterative basecalling workflow.** The per-site coverage was used as the metric to assess the efficacy of resolving compromised 3' and 5'-end basecalling. Within each coverage plot, the black curve mirrors the coverage of the counterparting plot.

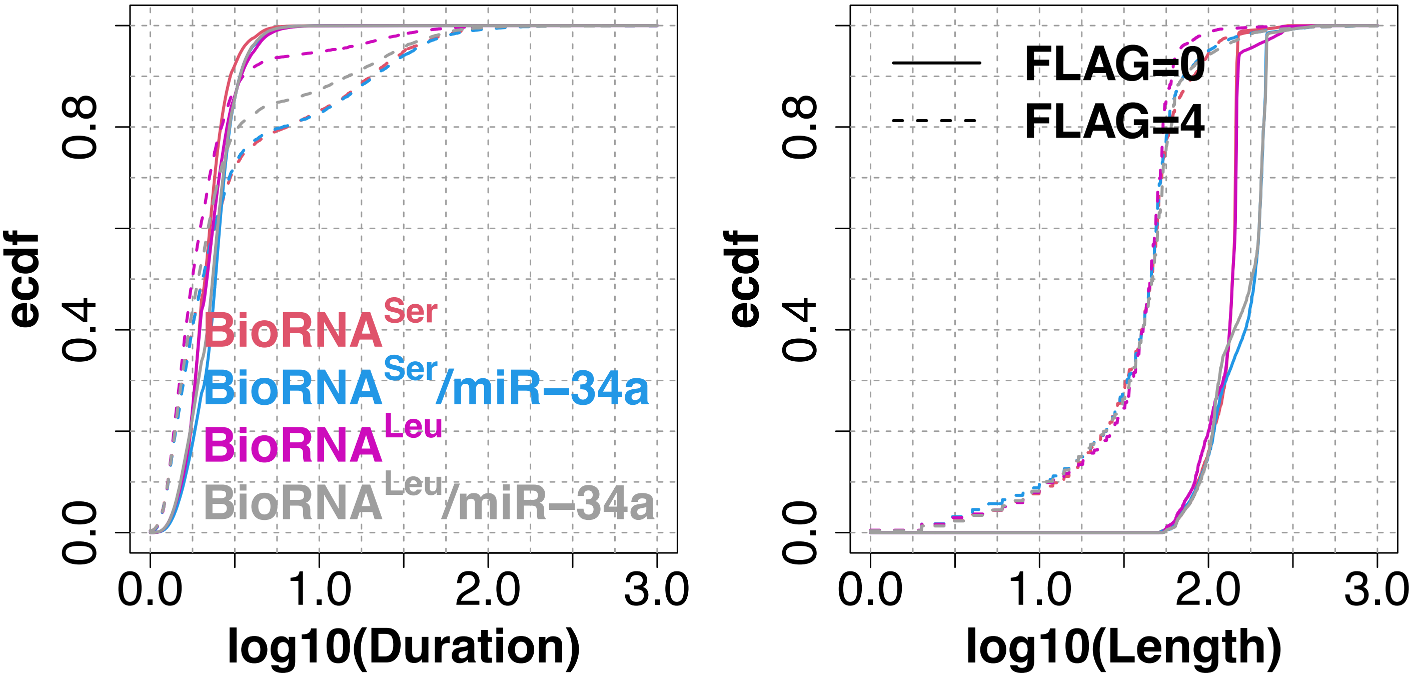

ecdf
0.0
0.4
0.8
0.0
1.0
2.0
3.0
log10(Duration)
BioRNA^Ser
BioRNA^Ser/miR−34a
BioRNA^Leu
BioRNA^Leu/miR−34a
ecdf
log10(Length)
FLAG=0
FLAG=4

**Figure S2. Distributions of dwell time and sequence length.** FLAG 0 and 4 denotes aligned and unaligned reads. Ecdf denotes empirical cumulative distribution function.

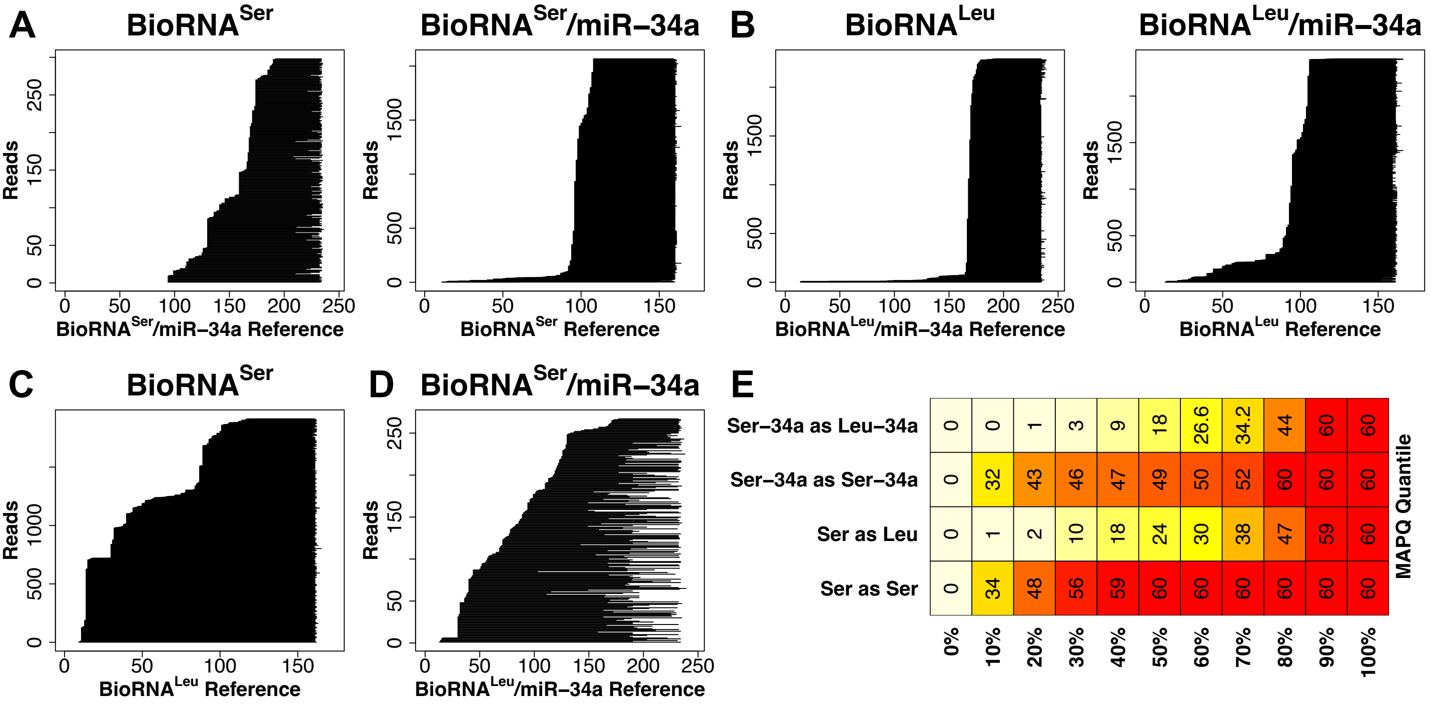
A
BioRNA^Ser
BioRNA^Ser/miR−34a
B
BioRNA^Leu
BioRNA^Leu/miR−34a
Reads
BioRNA^Ser/miR−34a Reference
BioRNA^Ser Reference
BioRNA^Leu/miR−34a Reference
BioRNA^Leu Reference
C
BioRNA^Ser
D
BioRNA^Ser/miR−34a
BioRNA^Leu Reference
BioRNA^Leu/miR−34a Reference
E
Ser−34a as Leu−34a
0 0 1 3 9 18 26.6 34.2 44 60 60
Ser−34a as Ser−34a
0 32 43 46 47 49 50 52 60 60 60
Ser as Leu
0 1 2 10 18 24 30 38 47 59 60
Ser as Ser
0 34 48 56 59 60 60 60 60 60 60
0% 10% 20% 30% 40% 50% 60% 70% 80% 90% 100%
MAPQ Quantile

**Figure S3. Mapping status of misaligned reads.** (A) Mapping positions of misaligned reads between BioRNA$^{Ser}$ and BioRNA$^{Ser}$/miR-34a. (B) Mapping positions of misaligned reads between BioRNA$^{Leu}$ and BioRNA$^{Leu}$/miR-34a. (C) Mapping positions of BioRNA$^{Ser}$ reads misaligned to BioRNA$^{Leu}$ reference. (D) Mapping positions of BioRNA$^{Ser}$/miR-34a reads misaligned to BioRNA$^{Leu}$/miR-34a reference. (E) MAPQ score distributions of (C) and (D) misalignments, as well as their corresponding correctly aligned counterparts.

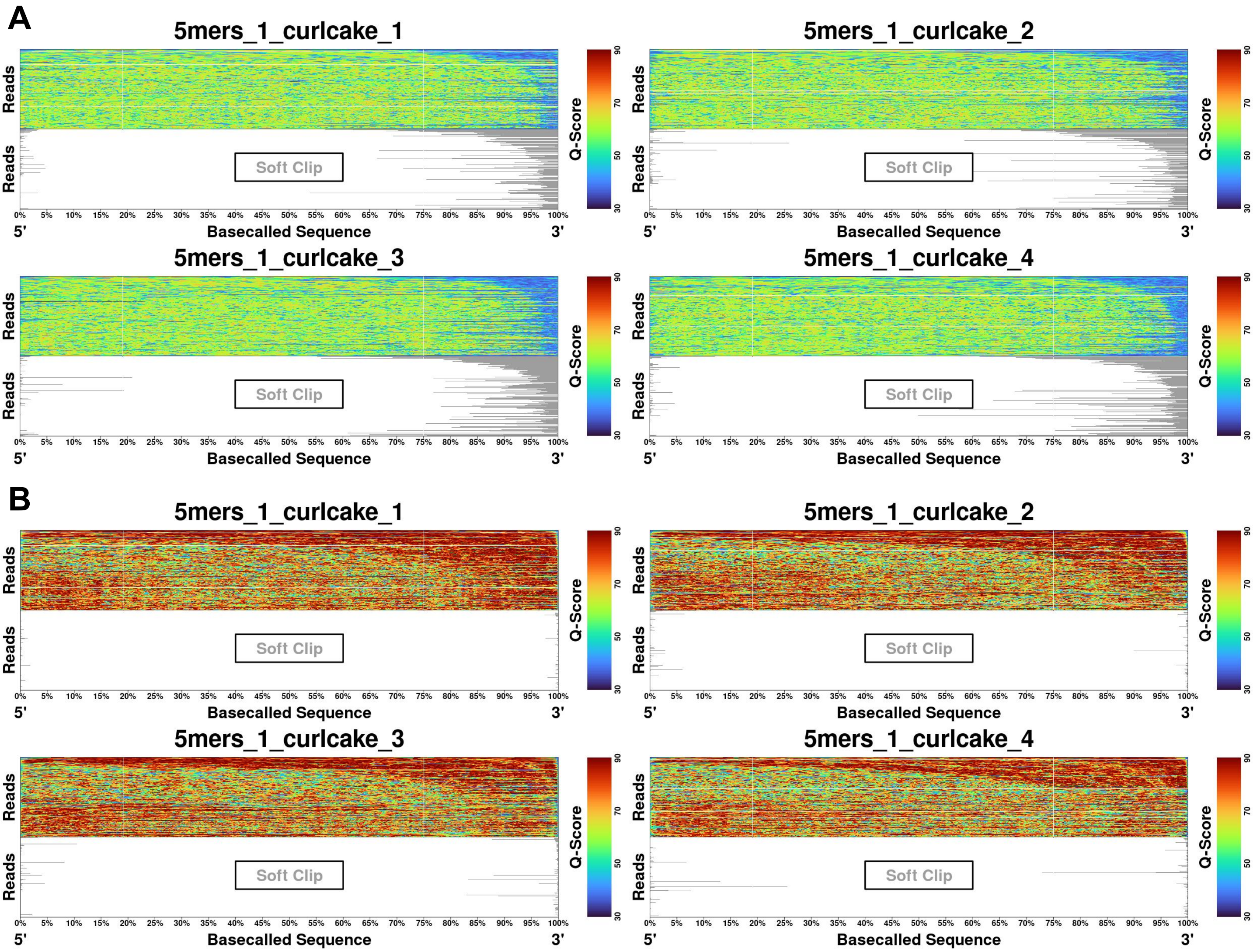

A
5mers_1_curlcake_1
5mers_1_curlcake_2
5mers_1_curlcake_3
5mers_1_curlcake_4
B
5mers_1_curlcake_1
5mers_1_curlcake_2
5mers_1_curlcake_3
5mers_1_curlcake_4
Reads
Soft Clip
Q-Score
30
50
70
90
0% 5% 10% 15% 20% 25% 30% 35% 40% 45% 50% 55% 60% 65% 70% 75% 80% 85% 90% 95% 100%
5'
Basecalled Sequence
3'

**Figure S4. Applying the 3-step sampling strategy to train long-read basecallers.** (A) Basecalling with the vanilla Bonito basecaller. (B) Basecalling with Bonito trained by the 3-step sampling strategy. Q-scores and soft-clipped regions were visualized along individual basecalled sequences. Sequences were normalized by lengths and visualized by length-percentages.

**Table S1. Sequences of BioRNAs and adapters.**

**Table S2. Sequences of synthesized RNA oligos.**
